# Toward an understanding of the small groups of thumb muscles that produce lateral pinch movement: application to surgical restoration of grasp following neurologic impairment

**DOI:** 10.1101/2024.09.04.611328

**Authors:** Cole D. Smith, Joseph D. Towles

**Author notes:** Corresponding Author: Joseph D Towles; Mailing Address: 500 College Ave Swarthmore, PA 19081.

## Abstract

**Purpose:** Tendon transfer surgeries that engage the flexor pollicis longus muscle (FPL) are commonly performed to enable lateral pinch grasp in persons with tetraplegia. Functional outcomes, however, have been mixed. This might be the case, in part, because FPL produces hyperflexion at the interphalangeal (IP) joint and radial deviation at the carpometacarpal (CMC) joint. Therefore, the goal of this simulation study was to investigate whether small groups of muscles could produce movement with less IP joint hyperflexion and CMC ab/adduction than FPL produces during lateral pinch movement.

**Methods:** We adapted a published, open-source computational musculoskeletal model of the hand to implement lateral pinch grasp. A forward-dynamics simulation approach was used to drive the thumb for 27 muscle groups being considered from an extended posture to a flexed posture to make contact with the side of the index finger. We calculated CMC ab/adduction deviation from the flexion-extension plane and IP joint flexion in the plane that all muscle groups produced, and compared those joint angle movements to those of FPL when it alone drove the thumb.

**Results:** Of the 27 simulations, three muscle groups, each consisting of three or four muscles, generated lower IP joint flexion and CMC ab/adduction than those of FPL.

**Conclusions:** This simulation work points to the potential of novel, multi-insertion site tendon transfer surgeries to out-perform the current standard of care to restore lateral pinch grasp following tetraplegia.

## INTRODUCTION

The quality of life of up to 135,000 persons around the world, who annually suffer a cervical spinal cord injury [1], is severely affected due to loss of hand function. Tendon transfer surgeries that engage the flexor pollicis longus (FPL) muscle are commonly performed to enable lateral pinch grasp [2]. Flexor pollicis longus is the most convenient muscle choice because it is the only thumb muscle that flexes all three joints and causes the least amount of radial-ulnar deviation [3]. Functional outcomes, however, have been mixed. That is, maximum, post-surgical pinch strength has differed by 10-fold among people and has been as low as tenths of a pound [4], [5]. This is possibly due to the oblique direction of FPL’s endpoint force that contributes to weak pinch force and a tendency for thumb-tip slip during grasp contact [6], [7], [8], [9]. From a joint movement perspective, FPL tends to cause hyperflexion of the interphalangeal (IP) joint due to the ratio of FPL’s flexion moment arms at the interphalangeal (IP), metacarpophalangeal (MP) and carpometacarpal (CMC) joints [3]. Interphalangeal joint hyperflexion diminishes the likelihood that the main portion of the thumbpad will make contact with the lateral aspect of the index finger. In this case, lateral pinch contact is not made at all, or is made between the very distal portion of the thumb-tip which, over time, induces callus formation on the thumb-tip. Stabilization of the IP joint by split-FPL tenodesis [10] or Steinman pin [11] has been used to reduce or eliminate, respectively, FPL’s tendency to hyperflex the IP joint and therefore enable thumbpad contact with the lateral index finger. From the perspective of the thumb-tip force that FPL produces, joint stabilization improves its oblique directional nature, but only to a slight degree [12]. Taken together, FPL’s endpoint force production and joint movement characteristics are less than ideal for the muscle to be the lone driver of the thumb.

More generally, mixed surgical outcomes to restore lateral pinch grasp after cervical spinal cord injury (SCI) or tetraplegia may exist because tendon transfer surgeries historically have not been designed with a biomechanics-based understanding of the endpoint velocities and endpoint forces that muscles generate during a grasping task. Previous studies, that have modeled multi-insertion site tendon transfer surgeries, have also shown that small groups of muscles produced more palmarly-directed forces (i.e., forces that better promote thumb-tip stability during pinch) than the current standard of care using FPL alone [7]. A multi-insertion site tendon transfer is one in which the non-paralyzed donor muscle is attached to multiple paralyzed recipient muscles (Fig. 1).

**Figure 1.**
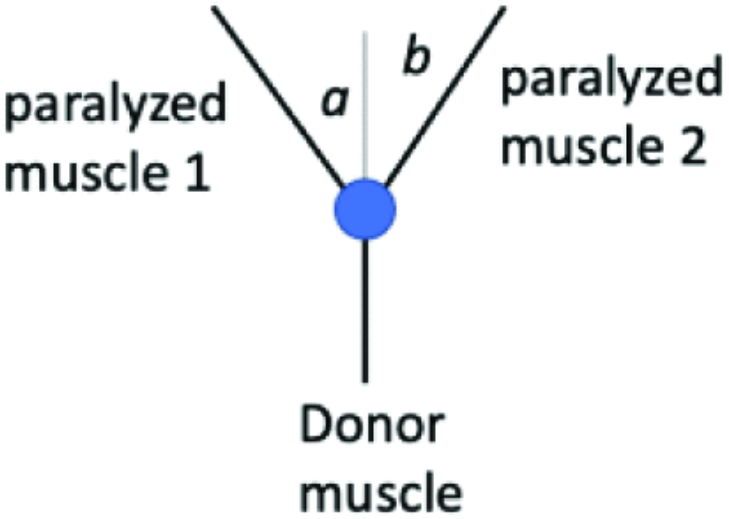
Multi-insertion site tendon transfer diagram. In this tendon transfer arrangement, the donor muscle is attached to multiple paralyzed recipient muscles. Two are pictured here. Donor muscle force is transferred to the tendons of paralyzed muscles 1 and 2. The proportion of force transferred to each tendon depends on the values of angles *a* and *b*

A preliminary execution of a mathematical model determined that only muscle combinations (e.g., multi-insertion site tendon transfers), and not individual muscles, have the capacity for the desired thumb-tip movement, and that these muscle combinations should include FPL. In a previous simulation study, we found that four groups of two muscles, ten groups of three muscles and 13 groups of four muscles produced well-directed endpoint forces during lateral pinch in the flexed thumb[7]. It is unknown, however, if these muscle combinations produce ideal lateral pinch movement from an extended to a flexed thumb posture.

The goal of this study was to investigate the joint movement characteristics of such muscle combinations in comparison to the current standard of care. Specifically, we wanted to know if they could produce movement of the thumb primarily in the flexion-extension (FE) plane as the thumb moved from an extended posture to a flexed posture making contact with the lateral aspect of the index finger to simulate lateral pinch movement. We hypothesized that at least one of the muscle combinations being considered would produce a more ideal movement pattern than the movement that FPL produces alone.

## MATERIALS AND METHODS

To investigate whether groups of thumb muscles could flex the thumb primarily in the FE plane from an extended thumb posture to a flexed one, we used a previously developed musculoskeletal model of the hand in the OpenSim framework. Specifically, our approach entailed adapting the model of the thumb [13] and implementing forward dynamic simulations of the thumb–driven by the muscle groups of interest–to facilitate lateral pinch from a wide-grip posture to a narrow one as the thumbpad made contact with the lateral aspect of the index finger.

### Adaptation of Musculoskeletal Model

The movement of the thumb-tip, driven by specific muscle combinations, was analyzed to determine if any sets of muscles were capable of producing ideal lateral pinch movement patterns of the thumb-tip throughout the FE plane. This was characterized by lower peak radial or ulnar deviation and IP joint flexion than that produced by FPL alone. These simulations were carried out using an adapted version of a previously developed hand and wrist model in OpenSim (Version 4.3) [13]. Briefly, the model consisted of flexion-extension and ab/aduction degrees of freedom (DOFs) at the CMC joint, a flexion-extension DOF at the MP joint, a flexion-extension DOF at the IP joint, and flexion-extension and ab/adduction DOFs at the wrist. The model also consisted of nine thumb muscles. Some thumb muscles have multiple names. For the purposes of comparing this study to the previous one [7] on which it is based, we will use thumb muscle names from that study and parenthetically note the alternate names used in the model [13] where it applies. The names of the nine thumb muscles were the flexor pollicis longus (FPL), the ulnar head of the flexor pollicis brevis (FPBu) (model: transverse head of the adductor pollicis), the radial head of the flexor pollicis brevis (FPBr) (model: flexor pollicis brevis), the adductor pollicis (ADP) (model: oblique head of the adductor pollicis), the opponens pollicis (OPP), the abductor pollicis brevis (APB), the abductor pollicis longus (APL), the extensor pollicis longus (EPL), and the extensor pollicis brevis (EPB). As to how the model was adapted, the model was adapted both in terms of the implementation of the elastic foundation used and the starting posture. Specifically, elastic foundation components were introduced as massless cylinders overlaying the thumb-tip, and the proximal, middle and distal phalanges of the index finger. These cylinders allowed the skin of the fingers to be approximated by massless rigid bodies and for contact force between these bodies to be calculated (Fig. 2). The thumb was then set to an extended position defined as 0 deg of flexion and abduction at the CMC joint, 0 deg of flexion at the MP joint and 0 deg of flexion at the IP joint (Fig. 2). As illustrated in Fig. 2, neutral flexion at the IP joint occurred when the long axes of the proximal and distal phalanges aligned and neutral flexion at the MP joint occurred when the long axes of the metacarpal and the proximal phalanx aligned. As in Smutz (1998)[3] (Paul Smutz et al., 1998), neutral flexion at the CMC joint occurs when the thumb rests on the side of the index finger in a lateral pinch posture. The other four digits were flexed to match the hand posture in a previous lateral pinch force study and the wrist was set to 0 deg of joint flexion and 0 deg of joint abduction (Fig. 2) [13], [14].

**Figure 2.**
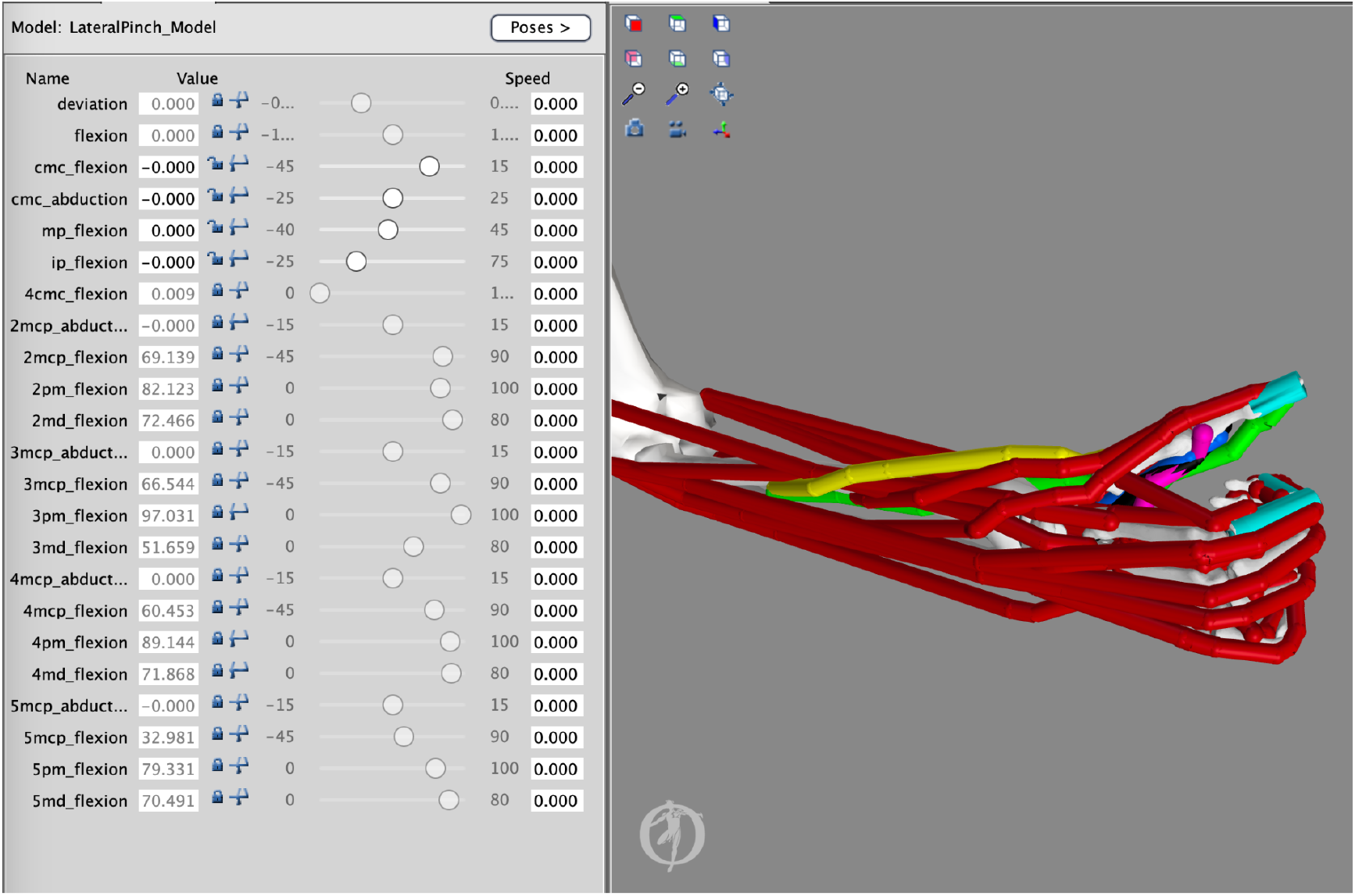
Adapted thumb model. The starting extended posture of the thumb was characterized by 0 deg of flexion and ab/adduction at the CMC joint, 0 deg of flexion at the MP joint and 0 degree of flexion at the IP joint. Massless bodies, attached to the thumb-tip and to the index finger, were used to detect contact between the thumb and index finger.

We then used this adapted model orientation to explore both muscle-group-driven and and FPL-driven thumb flexion from an extended thumb posture. The 27 muscle groups being investigated are consistent with the ones in a related previous study (Fig. 3) [7]. At the end of thumb flexion, the thumb made contact with the index finger as determined by the point at which the elastic foundation force deviated from zero.

**Figure 3.**
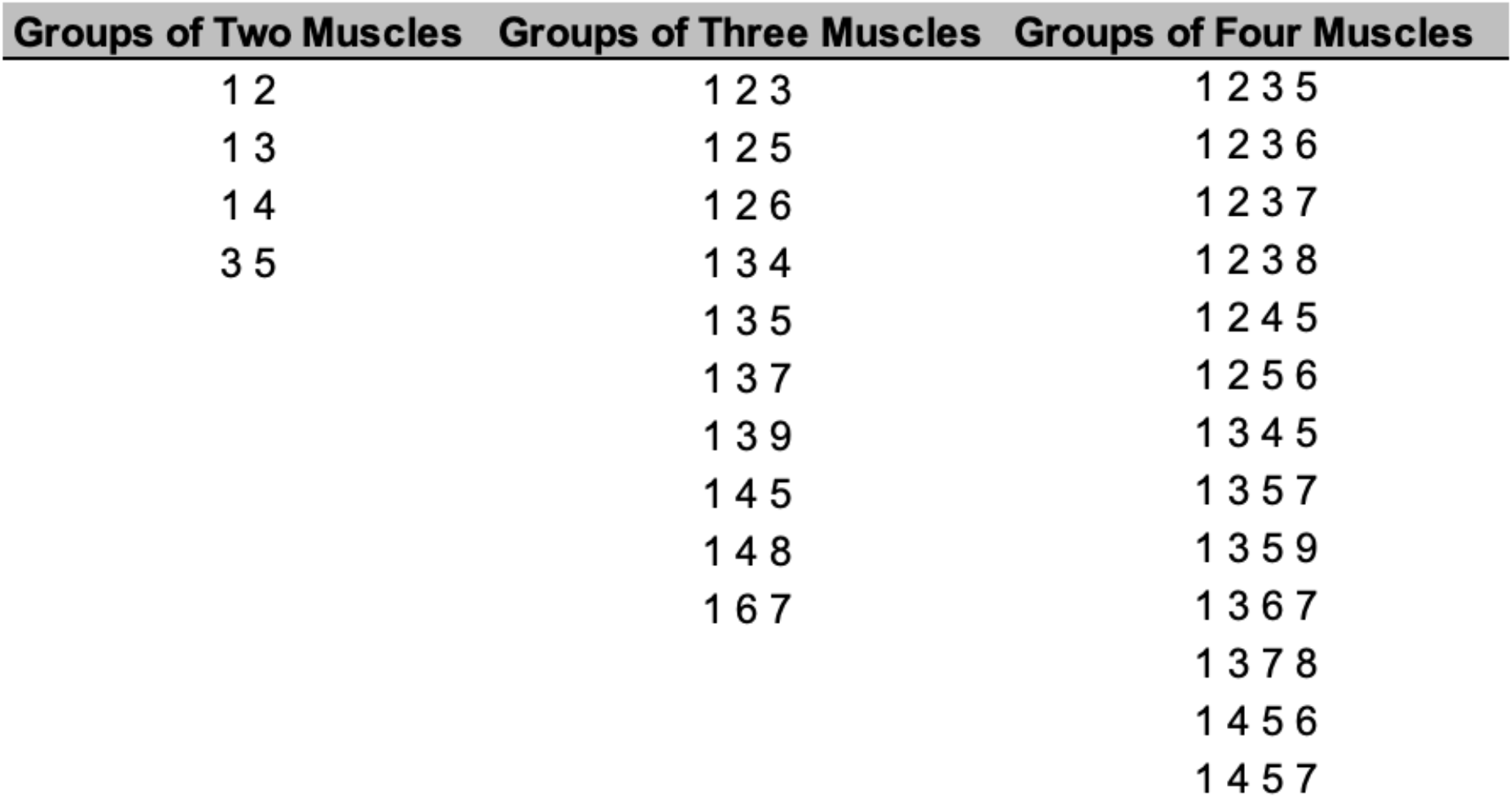
Thumb muscle groups. Muscle groups consisted of two, three or four muscles. Each number in a group represents a muscle in the following order: (1) FPL, (2) FPBr, (3) FPBu, (4) ADP, (5) APL, (6) EPL, (7) APB, (8) EPB, and (9) OPP.

### Model Simulation

To implement muscle-group-driven thumb flexion in OpenSim, the forward dynamics routine was used (Fig. 4). In the forwards dynamics routine, activation files were created for each muscle combination and for FPL such that only the muscles in the combination of interest or FPL were activated at 100% to actuate the thumb. When muscles were activated, they exerted their effect simultaneously on the thumb rather than in a coordinated fashion. From this, it also follows that muscles likely exerted different and various levels of muscle force throughout the range of motion (ROM) of lateral pinch movement. During the simulation, the finger joints were fixed in their flexed postures. The wrist joint was also fixed in its neutral posture. Twenty-eight simulations were carried out using OpenSim’s Forward Dynamics tool to investigate how FPL and each combination of muscles impacted thumb-tip movement during the execution of a lateral pinch grasp from an extended thumb posture to a flexed thumb posture. During this movement, we quantified IP joint flexion and CMC ab/adduction. Simulated thumb movements were integrated over 0.15 seconds, in line with other movement simulations used in the validation of the model [13], [14], allowing the thumb to make contact with the index finger and come to rest.

**Figure 4.**
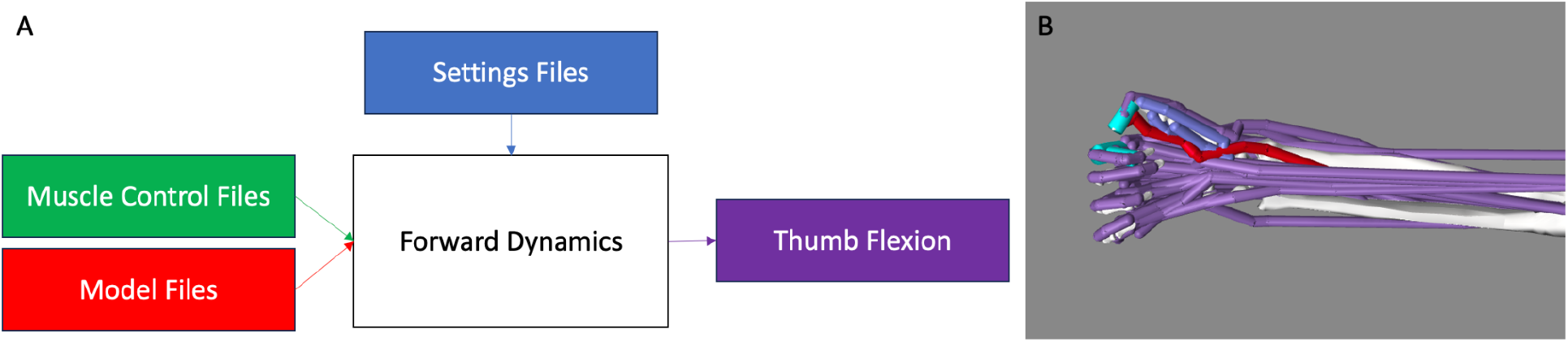
Overview of OpenSim forward dynamics framework. A) The adapted model (red) and .sto muscle control files (green) were loaded into the Forward Dynamics tool. A setting .xml file (blue) was then loaded to set the runtime of the model, set directories, and call for what results to be printed following completion of the simulation (purple). B) During forward dynamics, only activated muscles (red) drive movement and the remaining, non-activated muscles (purple) act passively. FPL was active in this case.

### Simulation Analysis

During lateral pinch movement, maximum IP joint flexion, maximum CMC abduction, and maximum CMC adduction were calculated relative to starting joint positions (using a MATLAB script). Carpometacarpal adduction and abduction values were combined to determine the range of motion outside the FE plane. Our analysis sought to determine muscle combinations that generated both less IP joint flexion and CMC ab/adduction compared to that produced by FPL alone. Maximum IP joint flexion and CMC joint range of motion were reported for each combination and ranked to determine which combinations minimized IP joint flexion, CMC joint ab/adduction, and both characteristics of the 28 simulations. Combinations that out-performed FPL in both respects were reported as combinations that could produce more favorable thumb-tip movement than FPL throughout the plane of lateral pinch movement.

## RESULTS

As we hypothesized, at least one muscle group produced less CMC ab/adduction and less IP flexion than those of FPL. Specifically, 11% or 3 of 27 muscle groups generated less IP joint flexion than FPL, i.e., less than 83 deg, and less CMC ab/adduction than FPL, i.e., less than 6.7 deg (Figs. 5,6). Specifically, FPL, FPBu, ADP and APL produced 53° of IP joint flexion; FPL, FPBr, FPBu and APB, 55°; and FPL, FPBu and APB, 65° (Figs. 5, 6). All 3 of the combinations contain FPL and FPBu.

**Figure 5.**
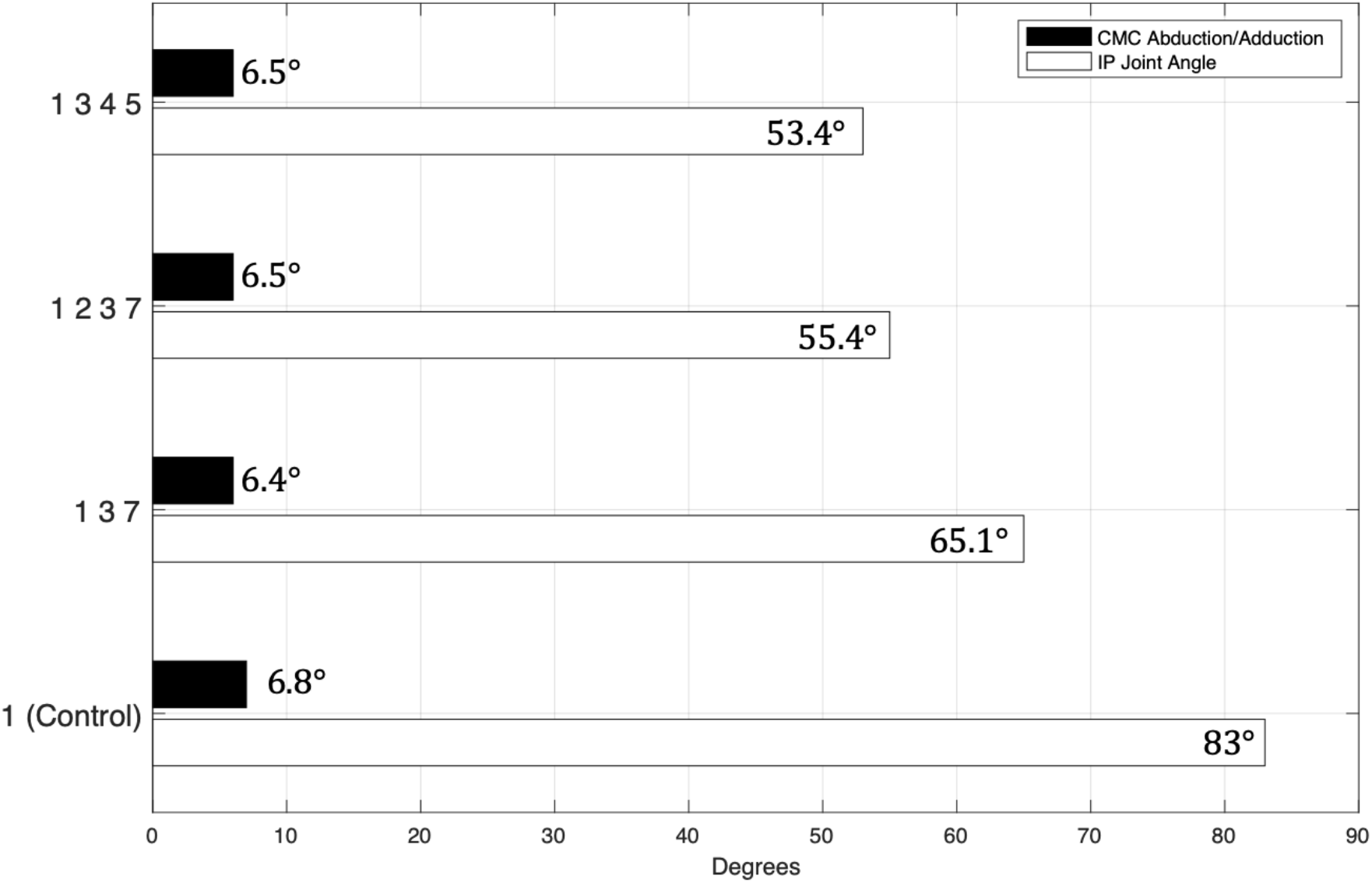
Combinations with improved joint movement characteristics. Muscle combinations which produced more favorable thumb-tip movement than FPL alone in both planes. The numbers represent the following muscles’ names: (1) FPL, (2) FPBr, (3) FPBu, (4) ADP, (5) APL, (6) EPL, (7) APB, (8) EPB, and (9) OPP.

**Figure 6.**
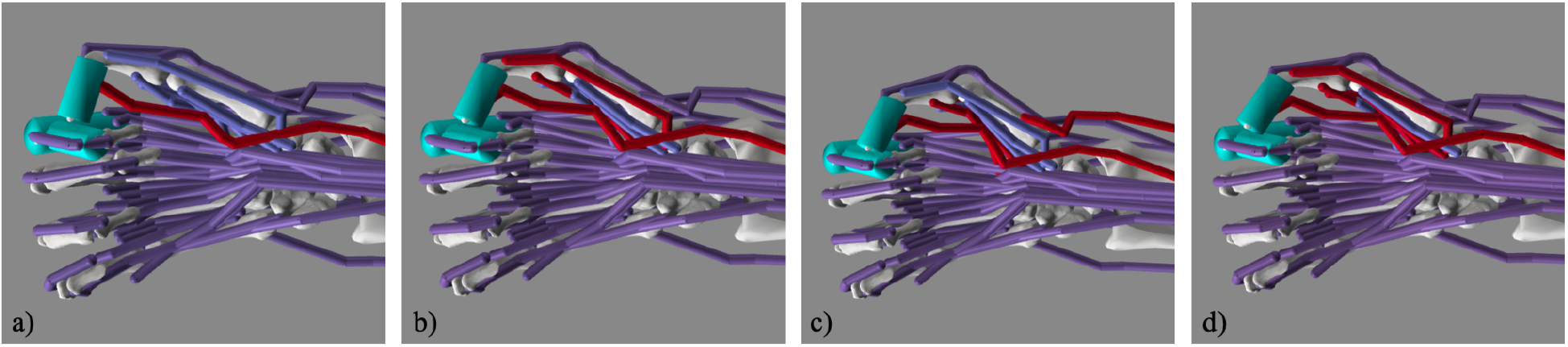
Simulated lateral pinch involving FPL only and various muscle groups. Images of the a) FPL b) FPL, FPBu, APB, c) FPL, FPBu, ADP, APL, and d) FPL, FPBr, FPBu, APB at contact in the simulation.

In all 28 simulations, the model achieved pinch, yielding an average maximum IP joint flexion of 70° and an average range of motion of 12° for abduction or adduction at the CMC joint. Flexion at the IP joint ranged from 41° (FPBu+APL) to 90° (EPL+FPL+APB), and ab/adduction at the CMC joint ranged from 6° (FPL+EPB+FPBu+APB) to 23° (EPL+FPL+FPBr+FPBu) (Fig 7).

**Figure 7.**
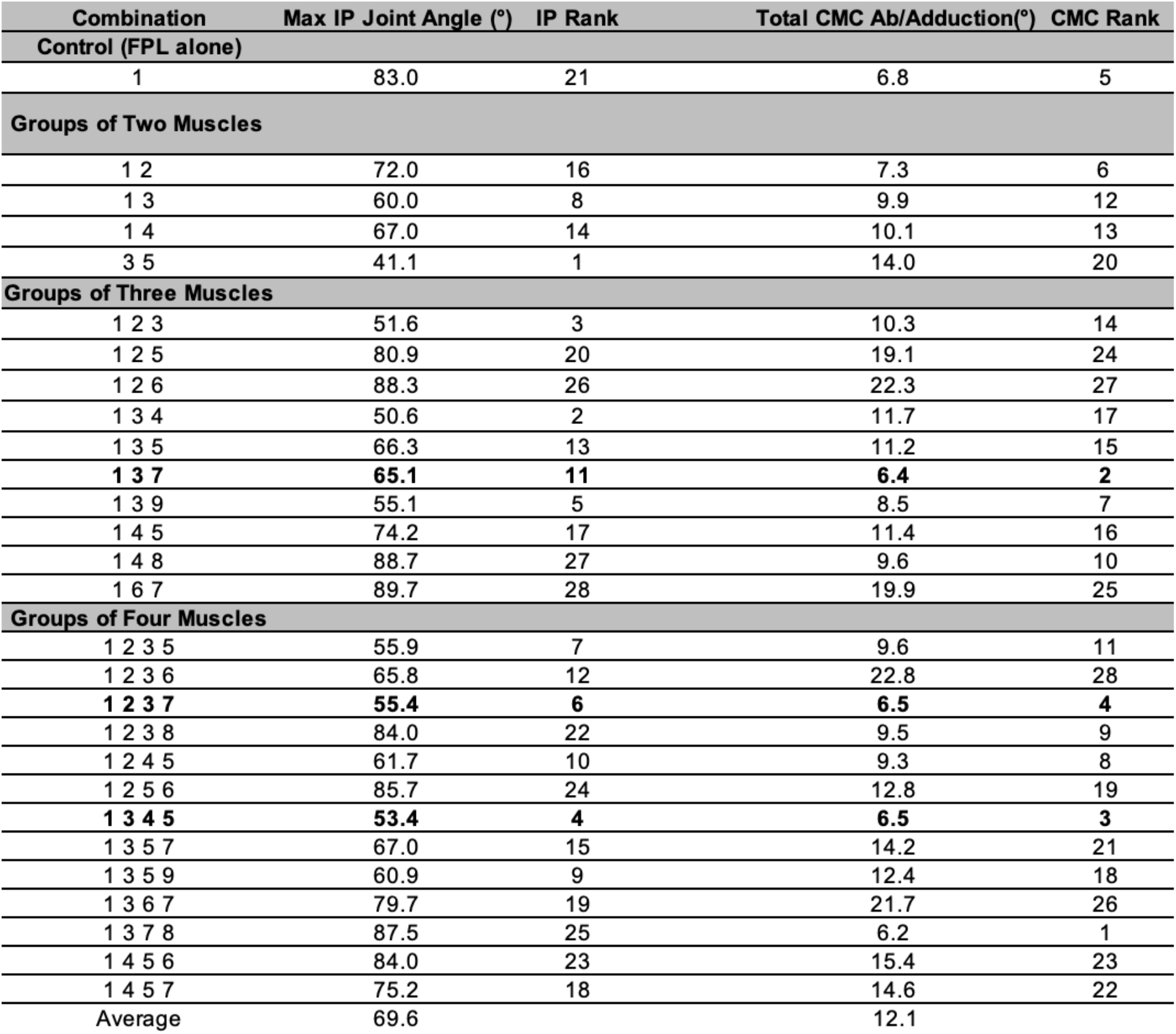
Forward dynamics simulation results. Maximum IP joint flexion, flexion rank, CMC joint range of motion, and range of motion rank are presented for the 28 simulations. The numbers represent the following muscles in the simulations: (1) FPL, (2) FPBr, (3) FPBu, (4) ADP, (5) APL, (6) EPL, (7) APB, (8) EPB, and (9) OPP.

Muscle groups out-performed FPL mostly in IP joint flexion and to a lesser degree in CMC joint ab/adduction. Of the 27 muscle-group simulations, 74% or 20 of 27 muscle groups flexed the IP joint less than FPL did alone and 15% or 4 of 27 muscle groups generated less CMC joint ab/adduction movement than FPL did alone. Both of these movements contributed to reduced IP joint hyperflexion and out-of-plane movement of the thumb during lateral pinch.

## DISCUSSION

The primary goal of this work was to show that, in a musculoskeletal model [13], combinations of small groups of muscles can generate improved thumb-tip movement patterns from an extended thumb posture to a flexed posture to achieve lateral pinch. This study was motivated by the need to find small groups of muscles that have both favorable endpoint force [7] and endpoint movement characteristics that out-perform those of FPL for the purpose of restoring lateral pinch grasp in persons following cervical SCI. As we hypothesized, multiple groups of muscles produced a more ideal lateral pinch movement pattern than the movement that FPL produces alone. Indeed, we found three such muscle groups that produced both less ab/adduction at the CMC joint and less flexion at the IP joint. Those muscle groups were (1) FPL, FPBu and APB; (2) FPL, FPBr, FPBu and APB; and (3) FPL, FPBu, ADP and APL. *To the best of our knowledge, this is the first simulation study to show the potential of small groups of thumb muscles to restore desirable lateral pinch movement*. The significance of this study is that it has the potential to improve lateral pinch grasp restorative surgeries in persons with cervical SCI because lateral pinch movement throughout its range of motion is being explicitly thought about from a biomechanical basis for the first time.

The results of the study can be explained by musculoskeletal mechanics. Experimental measurements of ab/adduction moment arm data for thumb muscles throughout the plane of FE do not exist. For reference, they exist for a single flexed thumb posture [3]. Notwithstanding, the findings that out-of-plane deviations at the CMC joint were comparable in three muscle groups (Fig. 5), during thumb flexion from an extended thumb posture to a flexed posture, could be explained by similar group-wide, average muscle length variations that are approximately the same as that of FPL. The 20 of 27 muscle groups that out-performed FPL (Fig. 7), by generating less IP joint flexion than FPL generated alone, primarily consisted of some combination of FPL plus uni- and/or biarticular muscles that flex the CMC joint only (uniarticular), and CMC and MP joints (biarticular) (Fig. 7). One such group was FPL combined with ADP and OPP, a biarticular muscle and a uniarticular muscle, respectively. In dynamic simulations, both uni- and biarticular muscles that flex the CMC only and flex both the CMC and MP joint can extend the IP joint. This non-intuitive action at the IP joint can occur because of complex dynamic coupling in a multi-joint musculoskeletal system. This may explain why these muscle groups produced less IP joint flexion than FPL produced alone.

Reduced IP joint hyperflexion and out-of-plane movement of the thumb during lateral pinch both can lead to a healthier and a higher-quality grasp than otherwise. Reduced hyperflexion of the IP joint reduces the risk of callus formation on the thumb-tip and creates greater thumb surface contact during grasping. The latter improves the quality of the contact during grasping because thumb pad surface torques can be passively resisted if need be [9]. Reduced out-of-plane movement has a greater potential for ideal contact between the thumb and the object being grasped than otherwise. By ideal contact, we mean that the thumb lands on the central portion of the grasped object. This thumb positioning improves the quality of grasp because the thumb would more likely than otherwise maintain contact with the object being grasped in the presence of any grasp disturbances.

The three muscle groups, which resulted in improved lateral pinch movement characteristics, contained both FPL and FPBu. The finding that FPBu is a common participant of these groups likely points to its broadly complementary endpoint velocity characteristics that support more favorable lateral pinch movement than FPL produces alone. This finding further supports the idea that intrinsic muscles have the potential to improve lateral pinch grasp characteristics (e.g., endpoint force and velocity directions) [7], [8]. As we stated early, we think that a small number of paralyzed muscles, being simultaneously driven by a donor muscle, can produce endpoint movement that is more favorably directed than FPL’s during lateral pinch grasp This would lead to improved grasp contact which has implications for grasp force production as well[9] (Murray et al., 2017). This idea that only muscle combinations (e.g., multi-insertion site tendon transfers), and not individual muscles, have the capacity for the desired thumb-tip movement, naturally follows from the understanding that multi-muscle control is required for accurate positioning of thumb and ultimately strong and stable grasps. A general tenet, however, of tendon transfer surgery is to attach one donor muscle to only one paralyzed recipient muscle [11]. That is to say, the tenet is to perform a single-insertion site tendon transfer. We believe that this tenet exists because of the difficulty of surgically planning, by inspection, the control of multiple recipient muscles by one donor muscle throughout the lateral pinch range of motion. Historically, computational musculoskeletal and cadaveric surgical simulation tools have not been used to plan lateral pinch tendon transfer surgeries. We believe that both can be brought to bear to investigate the potential of multi-insertion site tendon transfer surgeries.

As far as we know, this simulation study is the first to apply a musculoskeletal modeling approach to investigate the potential of small groups of thumb muscles to restore lateral pinch movement throughout the FE plane following cervical SCI. This work highlights the possibility of employing multi-insertion site tendon transfer surgeries to that end. Investigating the surgical feasibility of engaging multiple recipient muscles, including intrinsic muscles, in a tendon transfer surgery and determining how to implement the appropriate coordination of donor muscle force across recipient muscles are critical steps to consider.

## ACKNOWLEDGEMENTS

The authors would like to thank Dr. Daniel McFarland, at Guidance Engineers, for his assistance with the elastic foundation modeling.

## Statements and Declarations

The authors have no financial or non-financial interests to disclose that are directly or indirectly related to the work submitted for publication.

## Author contributions

All authors contributed to the study conception and design. Material preparation, data collection and analysis were performed by Cole Smith and Joseph Towles. The first draft of the manuscript was written by Cole Smith and Joseph Towles and all authors commented on previous versions of the manuscript. All authors read and approved the final manuscript.

